# Optogenetic induction of TAD boundary pairing promotes gene activity

**DOI:** 10.64898/2026.11.04.764356

**Authors:** Martin Stortz, Murali Palangat, Veer Keizer, Raj Chari, Daniel R. Larson, Tom Misteli

## Abstract

The genome folds inside the nucleus into distinct structures across multiple levels of organization. Topologically associating domains (TADs) are a ubiquitous architectural feature of genomes delineated by boundaries enriched in the CTCF protein and the cohesin complex. TADs are dynamic structures as their boundaries undergo continuous, short-lived pairing events. Whether the structural integrity of TADs contributes to regulation of the genes within the TAD is unclear. To directly test the structure-function relationship of TADs, we have developed OptoTet, an optogenetic tool to rapidly and directly manipulate chromatin contacts in a controlled fashion using light without the need for mutation or protein depletion. Using the TADs containing the *ERRFI1* or *MYC* gene as model systems, we show that OptoTet promotes TAD boundary looping. When applied to probing the structure-function relationship of TADs, light-induced pairing of TAD boundaries promoted transcriptional activity of *ERRFI1* as indicated by increased gene bursting. In contrast, bursting of *MYC* was insensitive to boundary pairing. Enhanced TAD boundary pairing also partially rescued the effect of depletion of cohesin components on *ERRF1* but not on *MYC*. Our results suggest that TADs boundary pairing dynamics can contribute to gene regulation in a gene-specific manner.

**GRAPHICAL ABSTRACT:** 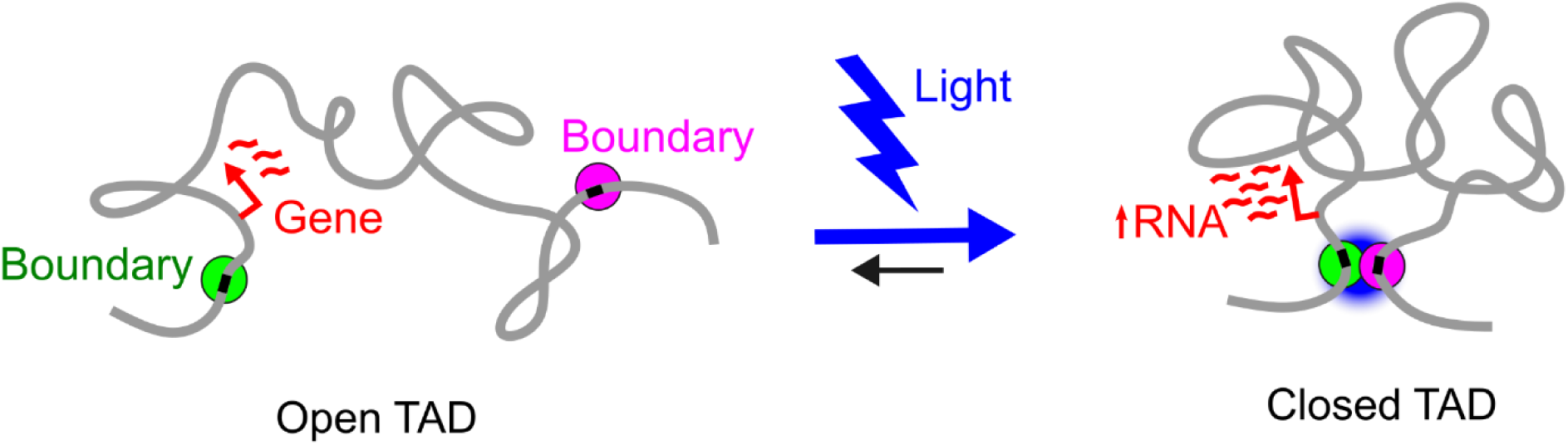

## INTRODUCTION

The human genome is folded inside the cell nucleus at multiple length scales, giving rise to distinct architectural features at different levels of organization. At the nanometer-scale, DNA wraps around core histones to form nucleosomes which assemble into the chromatin fiber [1]. The fiber then folds into loops of variable sizes which then form sub-micrometer topologically associating domains (TADs) [2]. Multiple TADs in turn coalesce into compartments based on their eu- or heterochromatin status [3]. At the highest level of organization at the micrometer-scale, chromosomes form and occupy preferred nuclear locations [4].

TADs have emerged as prominent and ubiquitous architectural features in the human genome [5]. A TAD is defined as a contiguous genomic region, typically ∼0.2-2 Mb in size, whose sequences undergoing preferential interactions among themselves rather than with other sequences [2, 5, 6]. TADs are formed by cohesin-driven loop-extrusion during which the cohesin ATPase activity pulls DNA through its ring-like complex to form a loop [7]. Extrusion arrests when the cohesin complex encounters the architectural protein CTCF bound to the TAD boundaries, forming a characteristic loop folded into a domain flanked by the boundaries [8–10].

Although TADs can be reliably mapped by genome-wide chromatin interaction profiling methods, the structure of TADs is highly variable between individual cells as indicated by single-cell imaging observations which found that any TAD is in a paired state only in a minor fraction of alleles in a population at any time [11–14]. The single-cell variability of TAD structure is in part due to dynamic motion of its boundaries as revealed by live cell-imaging which demonstrated that TAD boundaries are not stably associated with each other but undergo dynamic cycles of association and dissociation with persistence times of ∼20 min [15–17].

TADs have been proposed to play a functional role in gene regulation by favoring the physical interactions between gene promoters and regulatory elements, such as enhancers, within the same TAD and disfavoring inter-TAD interactions with regulatory sequences [18–21]. In support of such a role, disruption of TAD boundaries has been shown to affect the activity of several intra-TAD genes, including in the context of embryonic development and disease [21–26].

However, other observations suggest that TAD structure may not be a universal regulator of gene activity. In particular, functional enhancer-promoter interactions have been reported to occur across TAD boundaries [27–29], and genome-wide disruption of TADs only causes modest changes in gene expression [30–34]. Furthermore, the structural heterogeneity and dynamic nature of TADs complicate a potential regulatory role of TADs in the deterministic control of gene transcription [35–37].

A major challenge of uncovering the relationship between TAD structure and gene regulation is the limited availability of experimental tools to directly test structure-function relationships in chromatin. Mere correlations between changes in TAD architecture with changes in gene expression during differentiation, development, or disease may not reflect cause-effects relationships [31, 38–41]. Similarly, manipulation of TADs by experimental perturbations of architectural drivers such as mutation of sequence elements or depletion of architectural proteins, such as CTCF or cohesin components [21, 25, 28, 30, 33, 34], may lead to pleiotropic effects beyond those related to genome architecture [42, 43]. Furthermore, some of these perturbations occur at long time scales possibly leading to secondary indirect effects.

The limitations of current techniques have recently prompted the design and development of synthetic biology methods to manipulate genome architecture without the need to perturb key architectural proteins or sequences [44–49]. These methods have not yet been applied to manipulate TADs. We recently developed and demonstrated the effectiveness of OptoLoop [50], an optogenetic approach to manipulate chromatin contacts at short time scales and in a controlled fashion by applying light. OptoLoop consists of the fusion protein between a catalytically inactive Cas9 (dCas9) and CRY2, a protein from *Arabidopsis thaliana* which undergoes light-inducible clustering [51–53]. Given that OptoLoop allowed us to induce stable long-range chromatin contacts between repetitive DNA loci targeted with specific guide RNAs via dCas9 tethering [50], we reasoned that we could use a modified approach to manipulate TADs and examine the structure-function relationship of TAD boundary pairing on gene expression.

Here we describe ‘OptoTet’ a method to promote pairing of genomic loci in a controlled fashion using light. OptoTet is based on marking genome regions of interest with bacterial Tet operator (TetO) sequences, which are bound by its high-affinity and -specificity Tet repressor (TetR) ligand [54] fused to a variant of the CRY2 protein with enhanced clustering properties [50]. Using this approach, we demonstrate efficient induction of TAD boundary looping with light, favoring a closer, looped state of both the *ERRF1* and *MYC* TADs. We show that TAD boundary pairing leads to increased *ERRFI1* gene bursting both at the population and single allele levels. On the other hand, *MYC* gene bursting was not affected by TAD manipulation. Our results are consistent with the notion that TAD structure contributes to gene activation in a gene-specific manner.

## MATERIALS AND METHODS

### Cell lines

hTERT-RPE1-Cas9 is a telomerase-immortalized cell line from human retinal pigment epithelium (ATCC #CRL-4000) constitutively expressing CRISPR/Cas9 [55]. Lenti-X HEK-293T derives from human embryonic kidney (Takara Bio, Japan; cat. #632180,). All cell lines were cultured in Dulbecco’s modified Eagle’s medium with 110 mg/mL sodium pyruvate (Gibco, Waltham, MA, USA), supplemented with 10% fetal bovine serum (Gibco, Waltham, MA, USA) plus 100 IU/ml penicillin and 100 µg/ml streptomycin (Gibco, Waltham, MA, USA) at 37°C in a humidified atmosphere with 5% CO2. Cell lines were routinely checked for mycoplasma contamination.

### Plasmids

Plasmids were deposited in Addgene and are listed with their corresponding sources in Table S1. ERRFI1-5pTAD-96XTetO and MYC-5pTAD-96XTetO donor plasmids were generated in the Genome Modification Core (Frederick National Lab for Cancer Research) by assembling synthesized ∼800-1000 bp homology arms (Twist Biosciences, San Francisco, California, USA) with a 96XTetO construct obtained from Addgene plasmid #118713 [56]. ERRFI1-3pTAD-130XmTetO and MYC-3pTAD-130XmTetO donor plasmids were generated from a backbone generated in the Genome Modification Core and from a 72XmTetO construct generated as described [56]. mTetO repeats are made up each of a TetO half site and a mTetO half site (Supplementary Fig. S1). Briefly, two oligos carrying the mTetO sites and the restriction sites StyI, SpeI and BamHI were ligated together and then amplified with primers targeting the 5’-end of the first oligo and the 3’-end of the second oligo. The resulting PCR product corresponding to the length of the ligated oligo was gel purified and cloned into a TA cloning vector. The resulting plasmid was restricted with SpeI and BamHI and the StyI and BamHI digested fragment was ligated back into the plasmid. Since StyI and SpeI have compatible overhangs, ligation results in the loss of StyI and SpeI sites at the point of ligation and introduction of a new SpeI site downstream in the inserted fragment. The mTetO sites were multimerized by repeated digestion of the plasmid with SpeI/BamHI and reinsertion of the StyI/BamHI fragment to create a 72XmTetO repeat plasmid. The pHAGE_scTetR-3XGFP lentiviral plasmid was generated from the pHAGE_dCas9-3XGFP backbone (Addgene #64107[57]) and from a PCR-amplified single-chain TetR (scTetR) fragment from pCAG-TetR-Rad18UBD (Addgene #119010) [58]. scTetR binds as a single-chain dimer to wild-type TetO sites present in the 96XTetO arrays. The pHAGE_TetR-mTetR-3XmCh lentiviral plasmid was generated from the pHAGE_dCas9-3XmCherry backbone (Addgene #64108) [57] and from PCR-amplified TetR and mTetR fragments from the pHAGE_scTetR-3XGFP and pUBC_mutTetR-mCherry plasmid which was a kind gift from Cornelis Murre [59], respectively. The single-chain TetR-mTetR dimer binds to the TetO-mTetO sites in the 130XmTetO arrays [60]. pHAGE_scTetR-3XGFP-CRY2 and pHAGE_TetR-mTetR-3XmCh-CRY2 lentiviral plasmids were generated from pHAGE_scTetR-3XGFP and pHAGE_TetR-mTetR-3XmCh backbones, respectively, and from PCR-amplified CRY2 fragment from pCMV-CRY2hiclu-mCherry (Addgene #249577) [50]. psPAX2 (Addgene #12260) and pMD2.G (Addgene #12259) used for lentiviral transduction were obtained from Addgene.

### Generation of knock-in cell lines

CRISPR/knock-in integrations of TetO/mTetO repeats with their corresponding reagents and locations are listed in Tables S2, S3, and S4. Integrations were performed based on homology-directed repair [61]. Guide RNAs for targeting experiments were designed using sgRNA Scorer 2.0 [62] and their potency was assessed by measuring their CRISPR editing efficiency in HEK293T cells through Illumina sequencing as previously described [63]. Candidates with best editing efficiencies were subsequently used in knock-in experiments. First, integrations of 96XTetO at 5’ TAD boundary locations were performed. Target cells were subjected to reverse transfection by mixing 2 µg donor DNA plasmid, 30 pmol synthetic single-guide RNA (Synthego, Reedwood City, CA, USA) (Table S2), 12 µL Lipofectamine LTX (Thermo Fisher Scientific, Waltham, MA, USA), and 2.6 µL PLUS Reagent (Thermo Fisher Scientific) in serum-free media and then mixed with 7x10^5^ cells in a final volume of 2.5 mL growth media in a 6-cm dish. Target cells already constitutively express CRISPR/Cas9. After 3-5 days of recovery, selection of cells with genomic integrations of donor DNA was performed by incubation with 0.2 mg/mL Zeocin (InvivoGen, San Diego, CA, USA) for 1-2 weeks. Zeocin-resistant cells were seeded in 96-well plates at a one cell/well density using a MA900 Cell Sorter (Sony, Tokyo, Japan) for isolation of single-cell clones. Lysates from these clones were analyzed by PCR to test for integrations at the expected genomic locations (Table S3). For selected knocked-in clones, the integrity of the 96XTetO array was verified by PCR, and TetO spot visualization was tested by imaging after transient transfection with pHAGE_scTetR-3XGFP. A second round of integrations of 130XmTetO arrays at 3’ TAD boundary locations was performed using the selected clones with 96XTetO integrations at the 5’ TAD boundary as target cells as described above. In this case, 0.5 mg/mL G418 (InvivoGen) was used for selection. Two-color TetO/mTetO visualization and same allele integrations were verified by imaging after co-transfection of pHAGE_scTetR-3XGFP and pHAGE_TetR-mTetR-3XmCh. All integrations were confirmed to be heterozygous by PCR and/or imaging analysis.

### Generation of stable cell lines

Select knocked-in cell lines with same-allele integrations of 96XTetO and 130XmTetO at 5’ and 3’ TAD boundaries, respectively, were used for stable expression of TetR-GFP + mTetR-mCherry or TetR-GFP-CRY2 + mTetR-mCherry-CRY2 (Table S5). These stable cell lines were generated by lentiviral transduction in a sequential manner, i.e. first transduction of the GFP-fusion constructs and second, the mCherry-fusion constructs. Lentiviral particles were generated by transfecting a subconfluent 6-cm cell culture dish of Lenti-X HEK-293T cells with 1.7 µg psPAX2, 0.7 µg pMD2.G, 2.2 µg of lentiviral construct (Table S5), and 25 µl of 1 mg/ml polyethylenimine (ThermoFisher Scientific). After 48 h, supernatant with lentivirus was collected, filtered through a 0.45 µm filter, and concentrated overnight with Lenti-X Concentrator (Takara Bio, Shiga, Japan). Then, lentiviral particles were centrifuged (1500 g, 45 min) and resuspended in growth medium. Target cells were incubated with this lentiviral suspension and 10 µg/ml of Polybrene reagent (Sigma-Aldrich, Burlington, MA, USA) in subconfluent 24-well plates for 24 h. Transduced cell lines were sorted for mCherry- or GFP-positive cells with a Sony MA900 Cell Sorter to obtain polyclonal cell lines with variable expression levels. Single-clones were isolated by seeding polyclonal cell lines at very low density in a 15-cm dish and transferring single-colonies with PYREX cloning cylinders (Corning, Corning, NY, USA) after 2 weeks. Clones with different expression levels were selected for subsequent experiments.

### Light activation

CRY2 activation was performed by illuminating cells from above the plate with a 455-nm LED source (M455L3-C4, ThorLabs, Newton, NJ, USA) operated by a LED Driver DC2200 (ThorLabs, Newton, NJ, USA) as previously described [50]. Illumination was performed at pulses of 1 s every 10 s. Power was set at 5% at a current of 1000 mA, with a final power at sample of ∼1 mW/cm^2^. The duration of the illumination treatments was 3 h for DNA-FISH experiments and 4 h for RNA-FISH experiments. For controls, wells with cells kept in the dark were covered with aluminum foil. Cells were fixed with 4% paraformaldehyde (PFA) (Electron Microscopy Sciences, Hatfield, PA, USA) for 10 min at room temperature (RT) immediately after completion of the illumination protocol.

### CRISPR-knockout

Synthetic sgRNA kits commercially available for gene knockout from EditCo Bio (Redwood City, CA, USA) were used to knock out *RAD21* and *NIPBL* (Table S2). RNA was introduced in the target cells by reverse transfection in PhenoPlate 384-well microplate (Revvity, Waltham, MA, USA) by seeding 4x10^3^ cells with 1 pmol sgRNA and 0.2 µL Lipofectamine 2000 (Thermo Fisher Scientific) in a final volume of 50 µL growth media. After 72 h, cells were illuminated with a 455-nm LED and fixed with PFA as described above, followed by DNA- or RNA-FISH.

### DNA-FISH

For analyzing the effect of light alone, cells were seeded in a 384-well microplate at a density of 7x10^3^ cells per well. The next day, cells were illuminated with a 455-nm LED for 3 h as described above, followed by fixation with PFA. For CRISPR-knockout experiments, samples were prepared as described above. After three washes with PBS, cells were permeabilized with 0.5% Triton X-100 and 0.5% saponin in PBS (20 min, RT), washed twice with PBS, incubated with 0.1 N HCl (15 min, RT), and neutralized with 2X saline-sodium citrate (SSC) buffer (5 min, RT; #RGE-3240, K-D Medical, Columbia, MD, USA). Then, cells were pre-hybridized with 2X SSC, 50% formamide (overnight, 4°C). Fluorescently labeled BAC probes (Table S6) and commercial hybridization buffer were purchased from Empire Genomics (Buffalo, NY, USA). BAC probes were diluted in this buffer following the manufacturer’s instructions and added to the cells. Cells were then subjected to DNA denaturation (8 min, 85°C) followed immediately by hybridization (overnight, 37°C). The next day, hybridization mix was solubilized by adding 2X SSC (5 min, RT), cells were washed with 1X SSC (3x 5 min, 42°C) and with 0.1X SSC (3x 5 min, 42°C). Finally, cells were stained with 1 µg/ml DAPI (10 min, RT), washed and kept in PBS. The plate was subjected to high-throughput imaging.

### RNA-FISH

For analyzing the effect of light alone, cells were seeded in a 384-well microplate or in a Lab-Tek 8-well chambered coverglass (Thermo Fisher Scientific) at a density of 7x10^3^ cells or 3.5x10^4^ per well, respectively. The next day, cells were illuminated with a 455-nm LED for 4 h as detailed above, followed by fixation with PFA. For CRISPR-knockout experiments, samples were prepared at described above. For all *ERRFI1* RNA-FISH experiments, cells were maintained in medium with charcoal stripped FBS (Thermo Fisher Scientific) instead of regular FBS. After three washes with PBS, cells were permeabilized with 70% ethanol (2 h, RT), washed three times with PBS, and equilibrated with RNA-FISH wash buffer (2X SSC, 10% formamide) for 30 min at RT. RNA FISH probes (Stellaris) for RNAs were purchased as a pool of oligonucleotides labeled with Atto647 from LGC Biosearch Technologies (Hoddesdon, UK) as a commercial kit for *MYC* (cat. VSMF-2231-5) and custom designed for *ERRFI1* [64] (Table S7). Hybridization was performed overnight at 37°C with 10 nM probe, 2X SSC, 10% formamide, 10% dextran sulfate. Cells were then washed with RNA-FISH wash buffer (30 min, 37°C and 5 min, RT), PBS (2×5 min, RT) and stained with 1 μg/ml DAPI (10 min, RT). 384-well plates or 8-well plates were subjected to high-throughput imaging or confocal microscopy, respectively.

### Image acquisition

Fixed cell samples in 8-well chambered coverglass plates were imaged with a confocal laser scanning microscope LSM880 (Zeiss, Oberkochen, Germany), using an Alpha-Plan-Apochromat 100x/1.46 Oil DIC M27 Elyra objective. DAPI, GFP, mCherry, and Atto647 were excited with a 405 nm diode laser, an Ar laser at 488 nm, a 561 nm diode-pumped solid-state laser, and a 633 nm He-Ne laser, respectively. Fluorescence from these fluorophores was filtered in the 410-500 nm (DAPI), 500-544 nm (GFP), 588-633 nm (mCherry), and 652-708 nm (Atto647) ranges before detection with photomultipliers (DAPI and Atto647) or GaAsP spectral detectors (GFP and mCherry). Pinhole diameter was set to 0.9 µm. Single-plane images centered on TetO arrays of 64 x 64 pixels were acquired, with a pixel size of 66 nm. Signal was averaged 16 times with a final frame time of ∼10 s.

Fixed cell samples in 384-well plates were imaged with a CV8000 high-throughput spinning disk confocal microscope (Yokogawa, Tokyo, Japan) using a 60x water objective (NA=1.2). DAPI, GFP/’Green’, mCherry/’Red’/’Orange’, and Atto647 were excited with 405 nm, 488 nm, 561 nm and 640 nm lasers, respectively, and a 405/488/561/640 excitation dichroic mirror. Fluorescence was filtered through emission bandpass filters (445/45 nm, 525/50 nm, 600/37 nm, and 676/29 nm, respectively) and detected by two sCMOS cameras (2048 x 2048 pixels). Z-stacks spanning a range of 9 µm with 0.5 µm-intervals were acquired with a pixel size of 0.108 µm. Maximum z-projections, background correction, shading correction, and geometric corrections were performed on the fly by the proprietary Yokogawa software controlling the CV8000.

### Image segmentation

All imaging data were analyzed with the open-source High-Throughput Image Processing Software (HiTIPS) [65] using the computational resources of the NIH HPC Biowulf cluster (https://hpc.nih.gov). Nuclei from high-throughput images were segmented based on the DAPI channel using the deep learning model CellPose [66] implemented for GPUs. DNA- and RNA-FISH signals and TetO/mTetO arrays were segmented with the Gaussian Laplacian method as described previously [65]. Segmentation parameters were kept invariant for analyzing data from same experiments. The output of HiTIPS segmentation consisted of tables with single-nucleus and single-spot data, including size, shape, intensity, and spatial coordinates. Output segmentation data is publicly available on Figshare (https://figshare.com/projects/mistelilab_optotet/283089).

### Image analysis

Output data from HiTIPS was analyzed with the R software environment (R version 4.4.1, https://cran.r-project.org). Scripts are publicly available on Zenodo (https://doi.org/10.5281/zenodo.23508428).

For high-throughput DNA-FISH data, nuclei smaller than 70 µm^2^ or a solidity <0.95 were filtered out as a quality control step. Only nuclei with exactly two DNA-FISH signals for each channel were included in the analysis as hTERT-RPE1 are diploid cells. All possible distances between DNA-FISH signals from different channels in the same nucleus were computed using the spot centroids *x,y* coordinates. Minimum distances were considered as the single-allele distances between the targeted DNA loci by FISH as previously described [67].

For high-throughput RNA-FISH data, nuclei were filtered by size and solidity was determined as described above. The number of RNA signals per nucleus was computed. Only nuclei with 0, 1, or 2 RNA signals were included in the analysis. Burst fraction was calculated as the number of RNA signals over the number of alleles per cell as described [68].

For RNA-FISH coupled with TetO/mTetO imaging data, TetO/mTetO signals were segmented with a set of variable parameters since mTetO arrays were less prominent and more variable in shape/size resulting in some cases more than one mTetO spot per image. Images with no TetO or mTetO spots were discarded. The true TetO spot was selected as the one with highest intensity in the rare case of images with more than two TetO spots. All possible distances between TetO-mTetO spots in the same image were computed using the spot centroid *x,y* coordinates. Minimum distances were considered as the single-allele TetO-mTetO distances. An arbitrary distance threshold of 180 nm, similar to thresholds used in prior studies [12, 16] was selected to classify alleles as ‘proximal’ or ‘distal’ if they were below or above the threshold, respectively. Alleles with distances larger than 0.6 µm (*ERRFI1*) or 0.8 µm (*MYC*) were discarded from the analysis. Also, minimum TetO-RNA spot distances were computed for every image. Alleles with an RNA-FISH signal at a distance below a distance threshold (0.8 µm for *ERRFI1*; 1.5 µm for *MYC*) and with an integrated intensity above an intensity threshold (300 a.u.) were classified as ‘active’. Else, they were classified as ‘inactive’.

### Statistical analyses

All data was obtained from at least three independent experiments performed and measured on different days. Only relevant statistical comparisons are indicated in the figures with asterisks as *P<0.05, **P<0.01, ***P<0.001. Statistical analyses were done with GraphPad Prism. For multiple comparisons of means involving two factors, a two-way ANOVA was performed followed by Tukey’s, Bonferroni’s or Šídák’s multiple comparison tests (as indicated in the figure legends). For comparing means of two conditions, unpaired two-tailed t-tests were performed. For comparing fractions, Fisher’s exact tests and Breslow-Day tests were performed.

## RESULTS

### Engineering human TADs for single-allele visualization and manipulation

We sought to use the TetO/TetR platform to develop an experimental system to visualize TAD structure at the single-allele level and at the same time manipulate the interaction dynamics of TAD boundaries at short time scales in a controlled fashion. For this purpose, we engineered human retinal pigment epithelial (RPE1) cell lines by integrating TetO repeats at the boundaries of two model TADs. We selected *ERRFI1* and *MYC* as target TADs because of their distinct features (Fig. 1a). The *ERRFI1* TAD is a relatively small TAD (∼0.39 Mb) on human chromosome 1, with the *ERRFI1* gene residing only ∼65 kb away from the 5’ TAD boundary in addition to other genes throughout the domain (Fig. 1a). In contrast, *MYC* is located in a large TAD (∼2.8 Mb) on chromosome 8, is the only coding gene and resides in the middle of the TAD, far away from either boundary (Fig. 1a). Both TADs are defined by CTCF binding sites at the boundaries that function as anchors for the characteristic TAD loops observed in bulk contact maps (Fig. 1a, Supplementary Fig. S2).

**Fig. 1.**
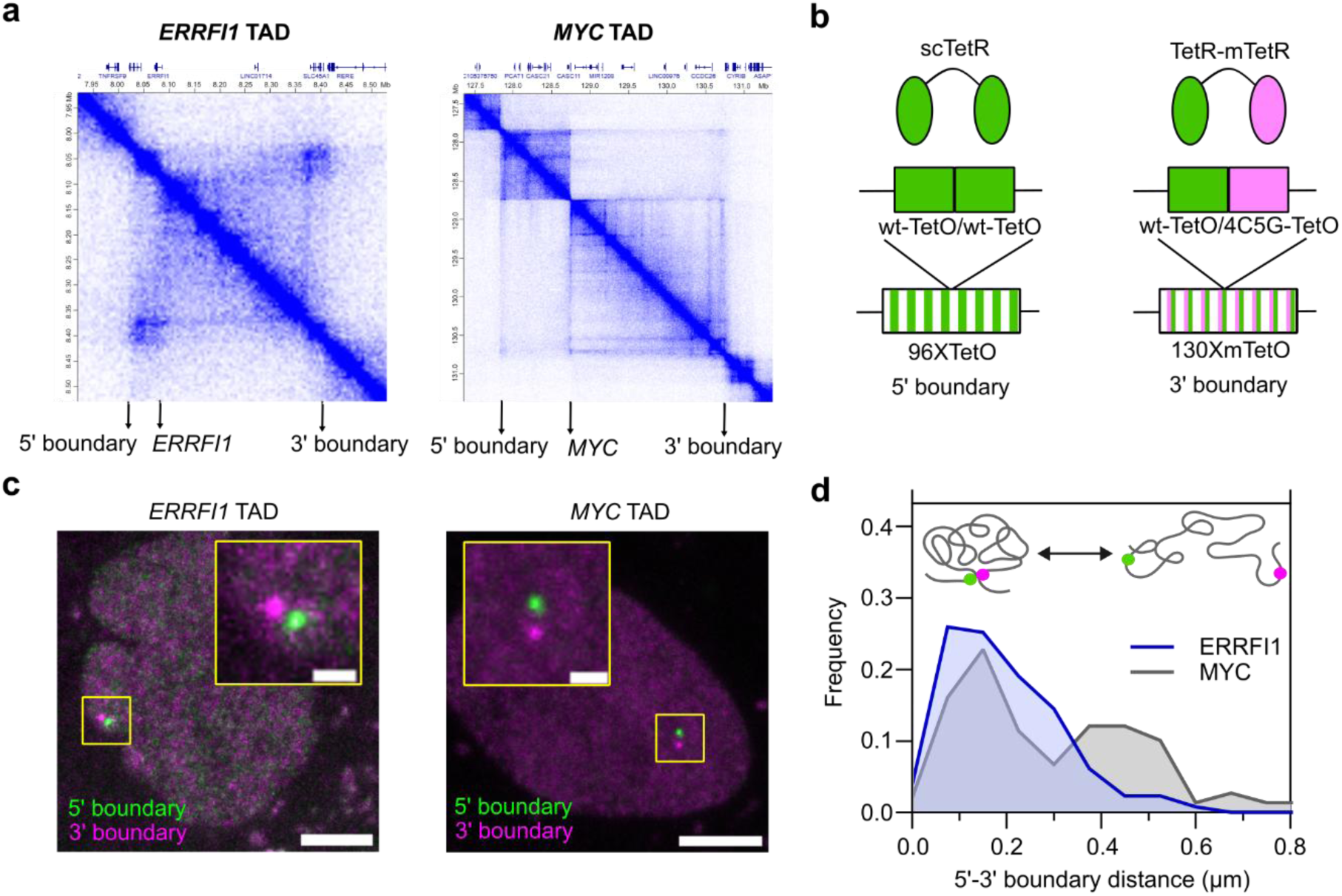
Engineering TADs for structure visualization and optogenetic manipulation. a) Hi-C contact maps for *ERRFI1* and *MYC* target TADs in hTERT-RPE1 cells were generated with Juicebox software [69] [70]. The location of the *ERRFI1* and *MYC* genes and their 5’ and 3’ TAD boundaries are indicated. Coordinates correspond to hg19 assembly. b) Schematic representation of Tet operator (TetO) integrations and Tet repressor (TetR) constructs used. c) Representative images of fixed RPE cells engineered for *ERRFI1* (left panel) and *MYC* TADs (right panel), expressing TetR-3XGFP and mTetR-3XmCherry. Insets show labeled TetO arrays (highlighted in yellow frames). Scale bars: 5 µm (large images), 1 µm (insets). d) Distributions of 5’-3’ TAD boundary distances in TetO-tagged RPE cell lines and expressing TetR-GFP-CRY2 and mTetR-mCherry-CRY2. 130-150 alleles were measured per sample from three independent experiments. Schematic representations of TAD structures are indicated for short and long boundary distances. Green and magenta dots represent TAD boundaries.

We integrated arrays of TetO repeats to create multiple binding sites for TetR (Fig. 1b), placing them ∼10 kb outside the TAD boundaries to avoid interference with loop-extrusion and native boundary function (Supplementary Fig. S2). To distinguish the 5’ and 3’ TAD boundaries, we integrated 96 wild-type TetO sequences at the 5’ boundary and 130 mutant TetO (mTetO) sequences at the 3’ boundary, which are recognized with high specificity by wild-type or mutant (mTetR) versions of TetR, respectively (Fig. 1b; see Methods for details)[59, 60, 71]. To prevent TetR/mTetR heterodimerization, we used single-chain TetR-TetR or TetR-mTetR dimers (Fig. 1b)[60, 72].

By transiently expressing TetR-GFP and mTetR-mCherry we were able to visualize the TetO arrays as prominent bright spots by confocal microscopy allowing simultaneous detection of the location of the 5’ and 3’ boundaries of the *ERRFI1* and *MYC* TADs (Fig. 1c). Consistent with the heterozygous nature of the engineered cell line, we detected only one pair of spots per cell (Fig. 1c). As expected, based on the dynamic nature of TADs, imaging of individual alleles revealed a range of spatial configurations of TAD boundaries as indicated by their center-to-center 2D distances in single-plane images, from boundaries completely overlapping to being separated by up to 0.5 µm (Fig. 1c-d; see Methods). These observations agree with previous distance measurements by FISH demonstrating heterogeneity in TAD boundary distances in a cell population [12, 13]. We surmise that overlapping boundaries likely represent a closed, looped state of the TAD, and separated boundaries represent a more relaxed, open state (Fig. 1d).

### Optogenetic induction of TAD boundary pairing

We sought to leverage the TetO/TetR system for TAD structure manipulation by optogenetics. Building on our recently described OptoLoop system to induce long-range chromatin interactions using light-induced clustering of CRY2 fusion proteins [50], we utilized TetR/mTetR as a DNA-targeting moiety for recruitment of TetR-CRY2 fusion protein to the TetO arrays at TAD boundaries (Fig. 2a). We reasoned that the activation of CRY2 clustering by light would stabilize the TAD boundary loop, manipulating the TAD structure by favoring its closed, looped configuration (Fig. 2a). We refer to this system based on targeting of CRY2 via TetO/TetR as ‘OptoTet’.

**Fig. 2.**
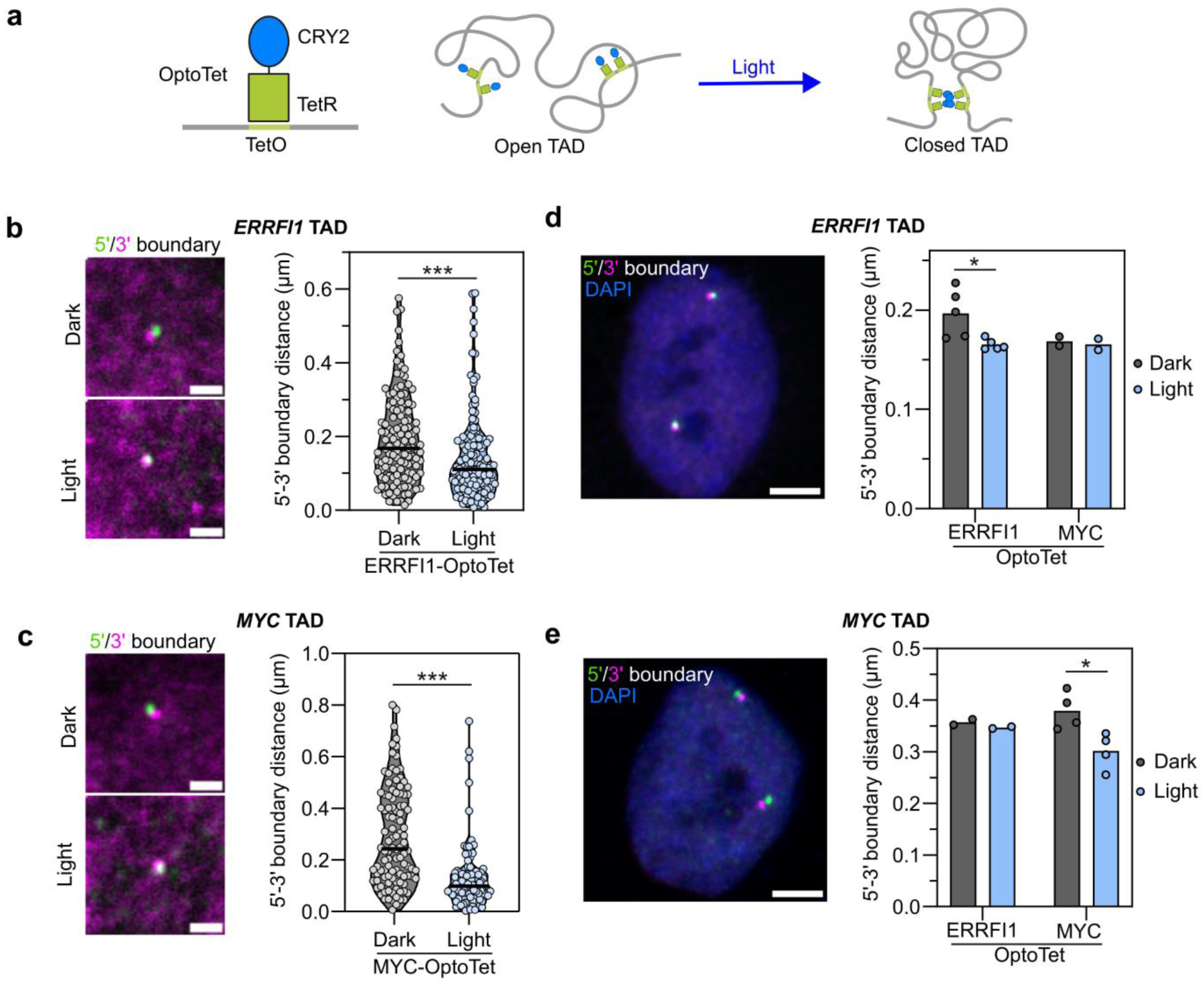
Optogenetic manipulation of TADs structure by OptoTet. a) Schematic illustration of OptoTet consisting of a fusion between TetR and the optogenetic protein CRY2, targeted to TetO arrays integrated into TAD boundaries. Light-induced CRY2 clustering stabilizes contacts between targeted TAD boundaries. b-c) Left panels, representative images of fixed RPE cells containing tagged 5’ and 3’ boundaries in (b) *ERRFI1* or (c) *MYC* TADs and expressing TetR-GFP-CRY2 and mTetR-mCherry-CRY2, kept in dark or illuminated with 455-nm light for 4 h (1 s pulses every 10 s). Scale bars: 1 µm. Right panels, 5’-3’ TAD boundary distances measured in each experimental condition. Each dot represents a single allele; black lines represent median distances. Data was collected from three independent experiments; 40-70 alleles were typically measured per experiment. d-e) Left panels, representative images of DNA-FISH performed with BAC probes targeting (d) *ERRFI1* and (e) *MYC* 5’-3’ TAD boundaries in RPE cells kept in the dark. Scale bars: 5 µm. Right panels, mean 5’-3’ TAD boundary distances measured from images of DNA-FISH performed with (d) *ERRFI1* or (e) *MYC* 5’-3’ TAD boundary probes in RPE cells engineered for *ERRFI1* or *MYC* TADs, expressing TetR-GFP-CRY2 and mTetR-mCherry-CRY2, kept in dark or illuminated with 455-nm light for 3 h (1 s pulses every 10 s). Each dot represents the mean of typically ∼1000-5000 alleles per condition per experiment. Bars represent the means of independent experiments. *p<0.05; **p<0.01; ***p<0.001 according to (b-c) t-tests, (d-e) two-way Anova followed by Šídák’s multiple comparisons tests. Only ‘Dark vs Light’ comparisons were analyzed.

We tested the ability of OptoTet to induce pairing of TAD boundaries in monoclonal RPE1 cells with TetO/mTetO tagged *ERRFI1* or *MYC* TADs and stably expressing low levels of TetR/mTetR-GFP/mCherry-CRY2. After illumination with 455-nm light using one-second pulses every 10 s for 4 h, we find a pronounced shift in the TAD distance distribution with a significant decrease in the mean 5’-3’ TAD boundary distance from 0.19 to 0.14 µm (p=0.0004) for the *ERRFI1* TAD (Fig. 2b) and from 0.29 to 0.13 µm (p<0.0001) for the *MYC* TAD (Fig. 2c) compared to control cells that were not illuminated. Similar boundary distances were observed in both TADs after light exposure, but as expected, the effect on boundary distance is smaller for *ERRF1* than for *MYC* given the smaller size of the *ERRF1* TAD with boundaries already closer in control conditions.

These results were confirmed by two-color, high-throughput DNA-FISH with bacterial artificial chromosome (BAC) probes targeting the *ERRF1* and *MYC* TAD boundaries (Fig. 2d-e). Consistent with the distance measurements based on TetO/TetR visualization, the mean 5’-3’ boundary distance decreased from 0.20 to 0.17 µm for *ERRFI1* TAD (p=0.022, Fig. 2d) and from 0.38 to 0.30 µm for *MYC* TAD (p=0.015, Fig. 2e) upon light-induced clustering of TetR/mTetR-CRY2. The absolute distance values are slightly larger when measured by FISH compared to direct visualization by TetO/TetR because BAC probes cover larger genomic regions thus providing a less accurate measurement of boundary distances. In control experiments, we verified the absence of off-target effects of light-activation by demonstrating that the mean 5’-3’ *ERRFI1* TAD boundary distance was unaltered in the MYC-OptoTet cell line (Fig. 2d) and vice-versa for *MYC* TAD in the ERRFI1-OptoTet cell line (Fig. 2e), indicating that OptoTet-driven proximity of TAD boundaries is locus-specific. Taken together, we have developed an optogenetic method to induce 5’-3’ TAD boundary contacts on short time scales, in a locus-specific and controlled manner.

### TAD boundary proximity promotes *ERRFI1* bursting

We next applied OptoTet to assess the effects of TAD structure manipulation on transcription bursting of the genes residing inside the engineered TADs (Fig. 3). For this purpose, we performed nascent RNA-FISH for *ERRFI1* (Fig. 3a) and *MYC* (Fig. 3b) and measured the fraction of active transcription sites (burst fraction) [68]. Light-induced TAD pairing increased the *ERRFI1* burst fraction in three monoclonal OptoTet cell lines expressing different amounts of the TetR-CRY2 fusion proteins, by typically 10-20% for all clones (Fig. 3a, Supplementary Fig. S3; p=0.025, 0.53, and 0.12 for clones #7, #11, and #4 respectively). *ERRFI1* bursting was not affected by light-exposure in parental cells not expressing any exogenous factors nor in cell lines expressing the Tet repressors alone without CRY2 moieties (Fig. 3a), eliminating the possibility of an artifactual effect of light exposure. These results suggest that induction of *ERRFI1* TAD boundary contacts promotes transcriptional activation of the *ERRFI1* gene. In contrast, while induction of boundary pairing was as efficient for *MYC* as for *ERRFI1*, *MYC* bursting was unaffected by light-induced pairing of *MYC* TAD boundaries (Fig. 3b, Supplementary Fig. S4). This difference may suggest gene-specific effects or may reflect differences in TAD properties such as the larger size of the *MYC* TAD.

**Fig. 3.**
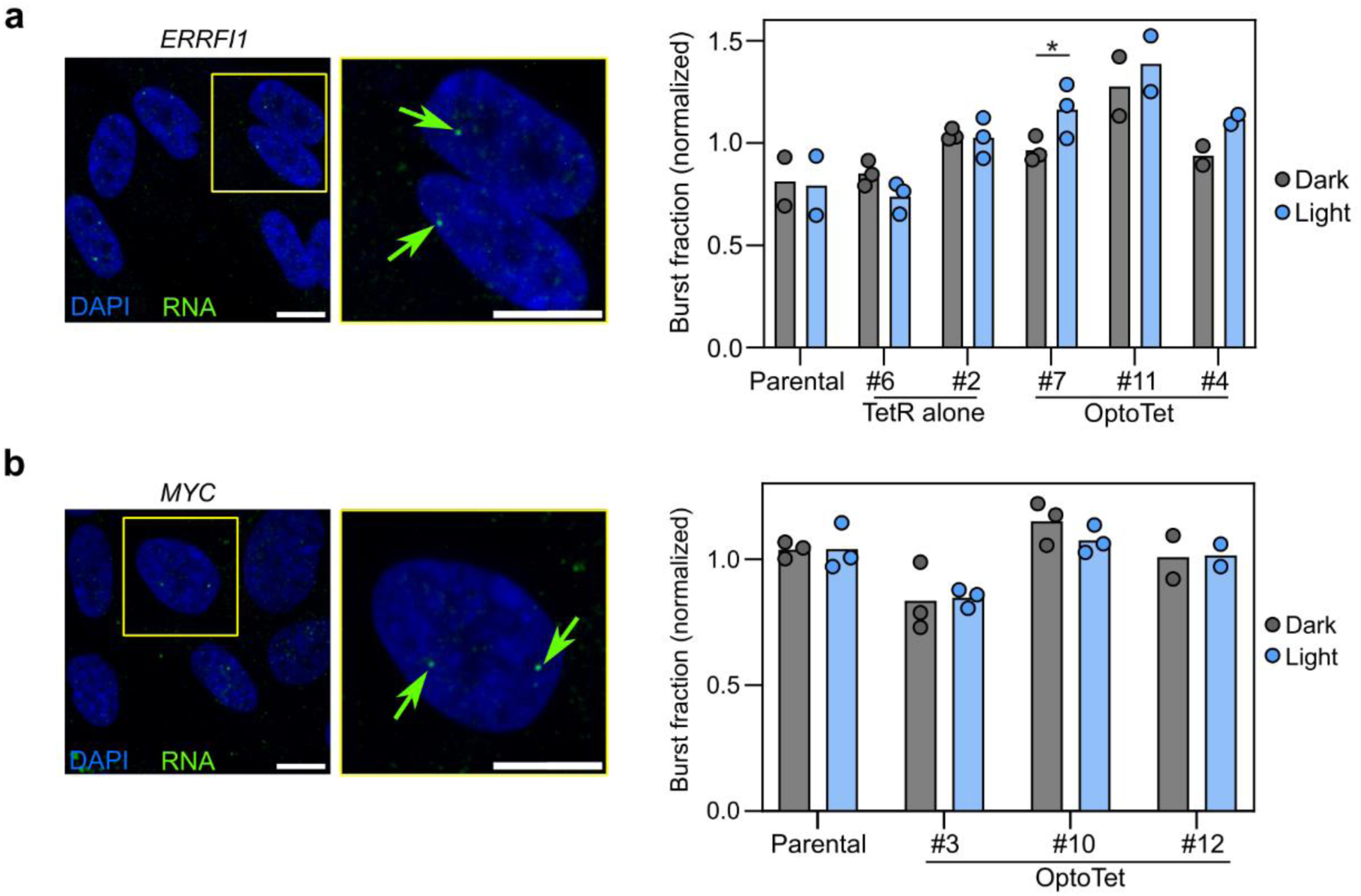
Effects of TADs structure manipulation on gene bursting. Left panels, representative images of nascent RNA-FISH performed with probes against (a) *ERRFI1* or (b) *MYC* pre-mRNA in RPE cells. Insets show cells (highlighted in yellow frames) with active transcription sites (indicated by green arrows). Scale bars: 10 µm. Right panels, burst fraction of (a) *ERRFI1* and (b) *MYC* genes measured from images of nascent RNA-FISH performed with the corresponding probes in RPE cells engineered for (a) *ERRFI1* or (b) *MYC* TADs, without any TetR constructs (Parental), or expressing TetR-GFP and mTetR-mCherry (TetR alone), or expressing TetR-GFP-CRY2 and mTetR-mCherry-CRY2 (OptoTet), kept in dark or illuminated with 455-nm light for 4 h (1 s pulses every 10 s). Clones with different expression levels were assessed (Supplementary Fig. S2-S3). Each dot represents the mean of typically ∼2000-6000 cells per condition per experiment. Bars represent the means of independent experiments. Burst fraction was normalized by the average burst fraction of all conditions for each experiment. *p<0.05; **p<0.01; ***p<0.001 according to two-way Anova followed by Šídák’s multiple comparisons tests. Only ‘Dark vs Light’ comparisons were analyzed for each cell line.

### Transcriptional activation of single alleles by TAD boundary pairing

Given that we observed an inverse correlation between mean *ERRFI1* TAD boundary distance and *ERRFI1* gene bursting frequency in the cell population after illumination, we wondered if this relationship existed at the single-allele level. Therefore, we sought to simultaneously analyze the gene activity status and the TAD structure of individual alleles. To do so, we coupled nascent RNA-FISH with 5’-3’ TetO array imaging and measured the 5’-3’ boundary distance of individual alleles, and at the same time classified each allele as active or inactive according to the presence or absence of RNA signal (Fig. 4a). We observed a correlation between the change in *ERRFI1* TAD boundary distance and allele activity. Upon light exposure, the mean boundary distance decreased from 0.21 to 0.12 µm for active alleles (p<0.0001), likely reflecting more persistent pairing of boundaries in response to light. No significant change in mean distance was observed for inactive alleles (0.18 to 0.16 µm, p=0.26, Fig. 4a). As an alternative, we computed the fractions of active and inactive alleles with a TAD distance of less or greater than 180 nm, a similar threshold as previously used to define TADs as open or paired [12]. We observed that, for unilluminated control cells, active alleles were roughly equally divided between close (<180 nm) and separated (>180 nm) *ERRFI1* TAD boundaries (Fig. 4a), in agreement with prior reports [13]. However, upon light induction, the fraction of active alleles with close TAD boundaries was considerably higher (55%) than with separate boundaries (32%, p=0.015) (Fig. 4a). Consistent with our observation that TAD boundary pairing does not affect *MYC* bursting (Fig. 3b), we observe for *MYC* a similar decrease in TAD boundary distance upon light for both active and inactive alleles and the fractions of active/inactive alleles between alleles with close and separated TAD boundaries remained similar (Fig. 4b). Overall, these data suggest that pairing of TAD boundaries can promote gene activation and do so in a gene-specific manner.

**Fig. 4.**
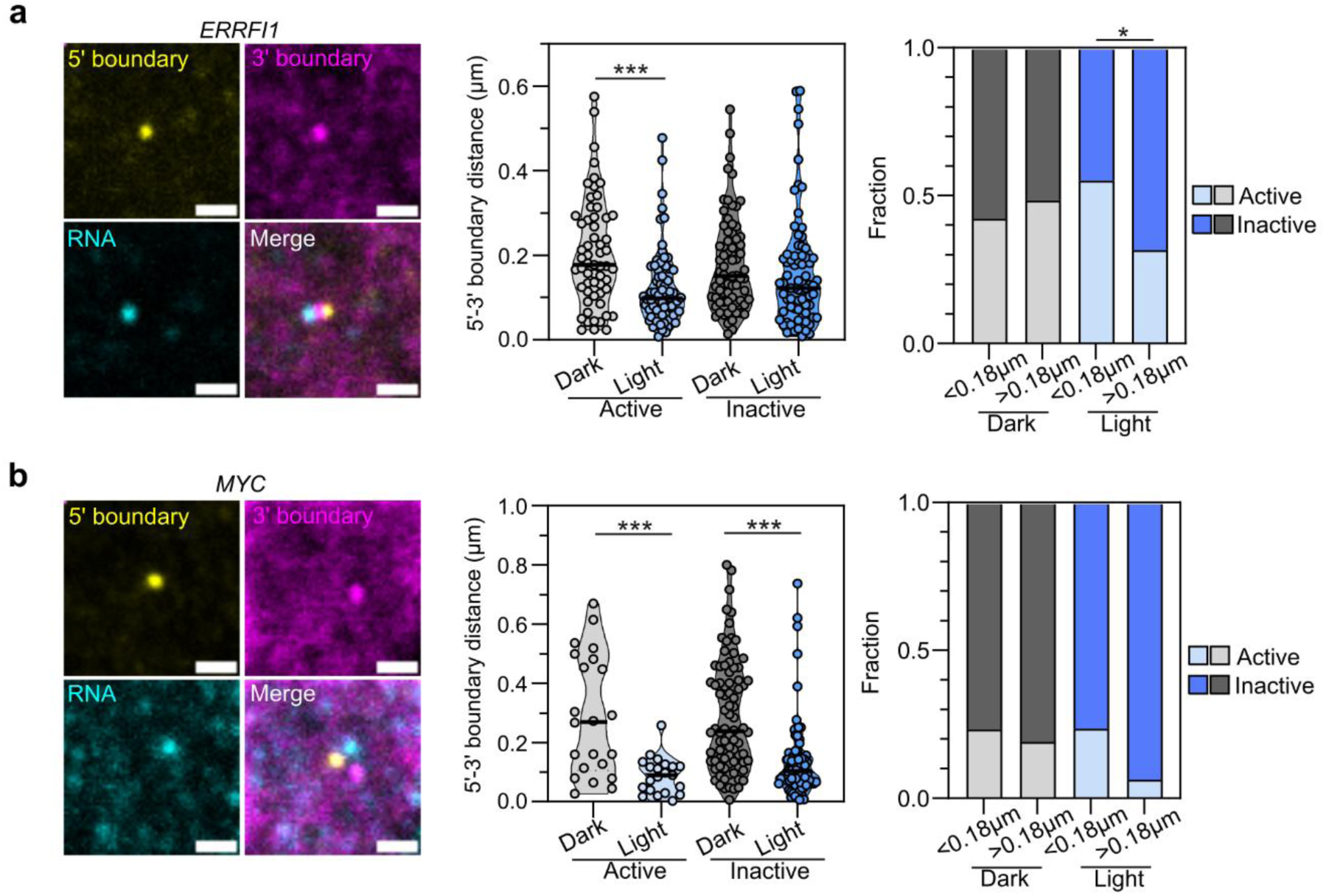
Link between TAD structure and gene bursting. Left panels, representative images of nascent RNA-FISH performed with probes against (a) *ERRFI1* and (b) *MYC* pre-mRNA on RPE cells engineered for (a) *ERRFI1* or (b) *MYC* TADs and expressing TetR-GFP-CRY2 and mTetR-mCherry-CRY2. 5’ and 3’ TetO arrays were visualized by GFP and mCherry channels, respectively. Scale bars: 1 µm. Middle panels, 5’-3’ TAD boundary distances measured from images obtained as described above for cells kept in the dark or illuminated with 455-nm light for 4 h (1 s pulses every 10 s). Alleles were classified as active if an RNA signal was within (a) 0.8 µm or (b) 1 µm. Each dot represents a single allele; black lines represent median distances. Right panels, fraction of active and inactive alleles obtained from experiments describe above. Alleles were classified according to their 5’-3’ boundary distance using a distance threshold of 180 nm. Data was collected from three independent experiments, typically 40-70 alleles per experiment were measured. *p<0.05; **p<0.01; ***p<0.001 according to two-way Anova followed by Tukey’s multiple comparisons tests, or Fisher’s exact tests. Only ‘Dark vs Light’, ‘Active vs Inactive’, and ‘<0.18 µm vs > 0.18 µm’ comparisons were analyzed.

### Forced TAD boundary pairing partially overcomes loss of cohesin components

Our results point to a functional role of TAD boundary pairing in gene regulation, at least for *ERRFI1*. We thus next asked whether the OptoTet-driven effects on TAD structure and on gene activation were dependent on the loop-extrusion process [18]. We leveraged that our RPE1 cell lines constitutively express the CRISPR/Cas9 endonuclease enabling analysis of the effect of TAD pairing after knockout (KO) of essential components of the cohesin complex which drives loop-extrusion [43] (Fig. 5).

**Fig. 5.**
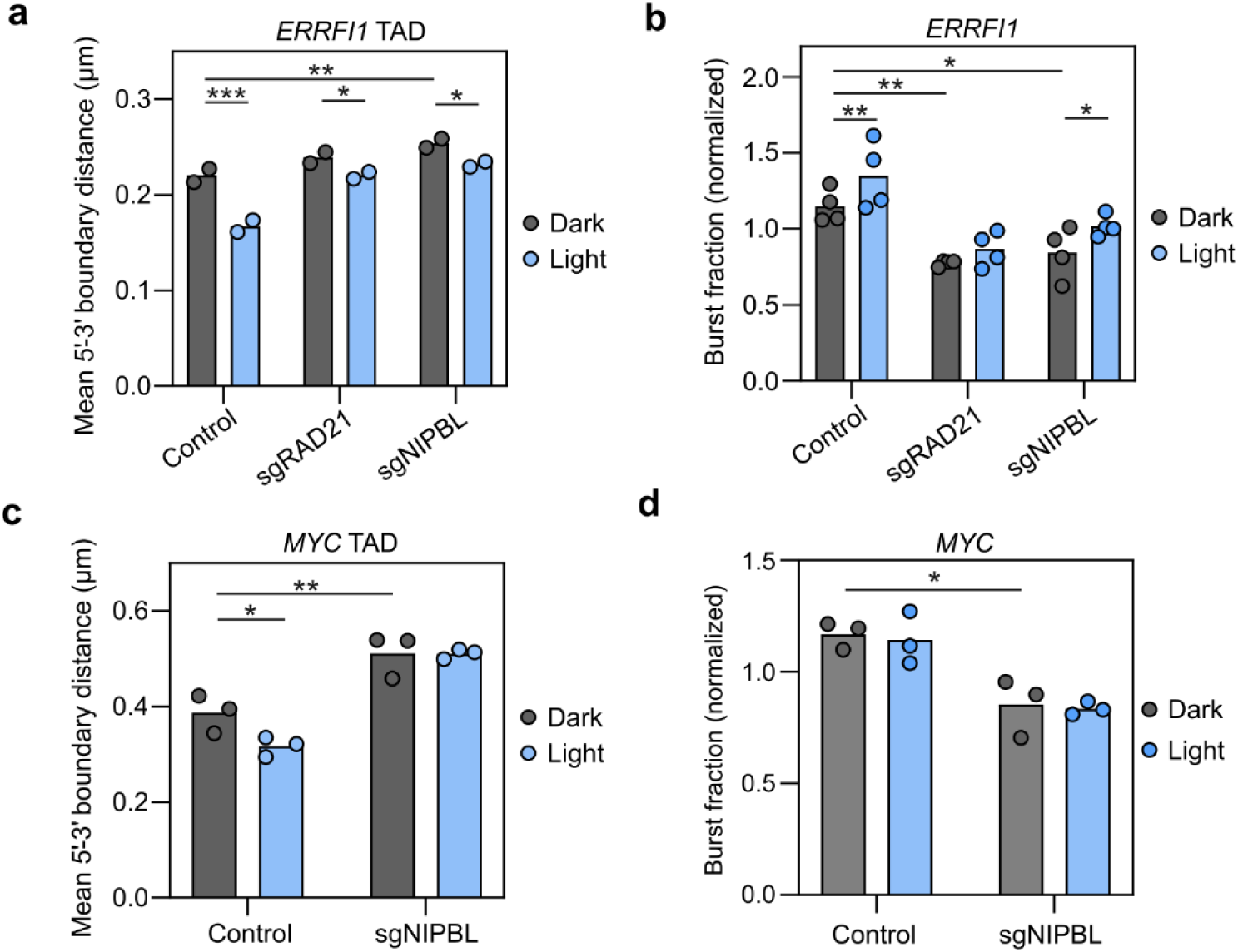
TAD structure manipulation and loss of cohesin components. RPE cells engineered for (a-b) *ERRFI1* or (c-d) *MYC* TADs and expressing TetR-GFP-CRY2 and mTetR-mCherry-CRY2 were subjected to CRISPR-knockout (KO) by transfection with sgRNAs targeting RAD21 or NIPBL. After 72 h, cells were kept in the dark or illuminated with 455-nm light for (a, c) 3h or (b, d) 4 h (1 s pulses every 10 s), then fixed. Control cells were not transfected with any sgRNA. (a, c) Mean 5’-3’ TAD boundary distances measured from images of DNA-FISH performed with BAC probes targeting (a) *ERRFI1* or (c) *MYC* TAD boundaries in cells as described above. Each dot represents the mean of typically ∼1000-5000 alleles per condition per experiment. Bars represent the means of independent experiments. (b, d) Burst fraction of (b) *ERRFI1* and (d) *MYC* genes measured from images of nascent RNA-FISH performed with the corresponding probes in cells as described above. Each dot represents the mean of typically ∼2000-6000 cells per condition per experiment. Bars represent the means of the independent experiments. Burst fraction was normalized by the average burst fraction of each experiment. *p<0.05; **p<0.01; ***p<0.001 according to two-way Anova followed by Šídák’s multiple comparisons tests. Only ‘Dark vs Light’ and ‘Control vs sgRNA’ comparisons were analyzed.

As previously reported, and in line with a role of cohesin in determining TAD structure [30], knockout of cohesin components RAD21 or NIPBL for 72 h, increased the mean 5’-3’ *ERRFI1* TAD boundary distance compared to control cells (from 0.22 to 0.25 µm, p=0.009) (Fig. 5a). Forced tethering via OptoTet after light exposure following KO partially rescued boundary separation (RAD21 KO, p=0.046; NIPBL KO, p=0.025). This extent of rescue was limited compared to the larger effect by light observed in control cells which express RAD21 and NIPBL () (Fig. 5a), suggesting that forced pairing of TADs can overcome some, but not all, of the inhibition of loop-extrusion due to loss of cohesin components.

We next analyzed the effects of light-induced TAD manipulation in the absence of loop extrusion factors on *ERRFI1* bursting (Fig. 5b). In line with prior observations [13], RAD21 or NIPBL KO reduced the *ERRFI1* burst fraction compared to control cells (Fig. 5b). Light treatment rescued some of the *ERRFI1* burst fraction (∼17% increase in control cells, p=0.0089; ∼12% increase in RAD21 KO cells, p=0.16; ∼21% increase in NIPBL KO, p=0.018; Fig. 5b).

In similar experiments for *MYC* (Fig. 5c-d), NIPBL KO led to a considerable increase in the mean 5’-3’ *MYC* TAD boundary distance (0.39 to 0.51 µm, p = 0.0016; Fig. 5c), whereas RAD21 KO resulted in reduced cell viability, precluding further analysis. In contrast to what we observed for *ERRFI1*, *MYC* TAD boundary looping was not rescued in NIPBL KO cells (Fig. 5c), likely because the larger *MYC* TAD depends more on loop-extrusion for boundary encounters. Similarly, in agreement with published data [73, 74], we also detected a significant decrease in the burst fraction of *MYC* gene upon NIPBL KO (Fig. 5d), but we did not observe any effects of forced TAD pairing on *MYC* bursting (Fig. 5d). These results support the notion that forced TAD boundary pairing can partially counteract the effect of loss of cohesin components on TAD structure and activity of some genes.

## DISCUSSION

We have developed OptoTet, a synthetic biology tool based on optogenetics to induce chromatin interactions between engineered genomic regions in a controlled fashion. We applied OptoTet to manipulate the structure of TADs in human cells to probe the relationship of TAD architecture and gene activity. We demonstrate efficient light-induced pairing of the boundaries of the *ERRFI1* and *MYC* TADs. Increased TAD boundary contacts were associated with an elevation in gene bursting of *ERRFI1*, but not of *MYC*, suggesting a gene-specific link between TAD structure and gene regulation.

TADs are highly dynamic structures [15–17]. Individual TADs in a single cell remain in a fully looped state with their boundaries in proximity for only a relatively short period of time with boundary pairing times estimated to be on the order of 20 min [15, 16]. The rationale for OptoTet to manipulate 3D genome organization was to enhance and stabilize interactions between genome regions marked by the TetR/TetO system via the light-induced clustering of CRY2 [50]. We indeed find that TAD boundaries for both model genes were more frequently in proximity in a cell population upon light illumination. The most likely scenario for the observed increased proximity of TAD boundaries is that their spontaneously occuring interactions are stabilized leading to prolonged persistence times. It seems less likely that OptoTet modifies the frequency of random encounters between targeted loci, for example by formation of large TetR-CRY2 aggregates, since the 3D search process is controlled by biophysical and biochemical constraints such as chromatin polymer mechanics, molecular motors, and architectural chromatin proteins [17, 59, 75, 76]. In line with that view, the ability to induce TAD boundary proximity with OptoTet was reduced in the absence of the cohesin components RAD21 or NIPBL, probably because the number of boundary encounters is reduced when loop extrusion is inhibited. Further support for this interpretation is our observation that the effects of cohesin knockout on OptoTet performance were more dramatic for the large *MYC* TAD with boundaries located 2.8 Mb away from each other, thus strongly depending on loop extrusion for encounters, whereas knockout of NIPBL or RAD21 only partially inhibited OptoTet manipulation of the *ERRFI1* TAD, whose boundaries have a higher encounter probability due to its smaller size. These observations support the notion that OptoTet does not operate by inducing the frequency of encounters between boundaries but by making natural encounters more stable via CRY2 multimerization upon light induction.

Our results point to a nuanced and gene-specific link between TAD boundary pairing and gene activation. The observation that the expression of *MYC* was unrelated to the pairing frequency of its TAD boundaries demonstrates that TAD architecture is not an obligatory determinant of gene activity. This conclusion is in line with prior observations by FISH which found that TAD architecture and gene activity are uncoupled [13]. The observed effect on gene activity of increased boundary pairing in the *ERRFI1* TAD is, however, consistent with the traditional model of TADs activating gene transcription via a closed, looped configuration by promoting productive intra-TAD contacts between genes and regulatory elements [18, 36, 37]. In the case of *ERRFI1* this regulatory effect may be due to the gene’s relatively close proximity to the 5’ boundary, with the forced pairing of *ERRFI1*

TAD boundaries increasing the chances of persistent interactions between the gene promoter and enhancers located near the 3’ boundary. It is unlikely that the effect on *ERRFI1* reflects altered chromatin behavior due to TetR-CRY2 tethering since we observed no change in bursting behavior relative to parental cells. Indeed, the observed susceptibility of boundary proximal *ERRFI1* to TAD structure is consistent with evidence showing that disruption of TADs preferentially affects the transcription of genes located near boundaries [77]. This property would also explain why the expression of *MYC* was not affected by forced *MYC* TAD boundary pairing, since the gene is at the center of the TAD, more than 1 Mb away from the closest boundary.

A major advantage of OptoTet compared to traditional approaches is that it allows us to directly manipulate TAD structure rather than rely on correlative observations. For example, using OptoTet in the absence of illumination, we reproduced earlier observations using simultaneous RNA/DNA FISH analysis of *ERRFI1*, in which we found no correlation between TAD boundary pairing and gene bursting [13]. However, we extended those findings by demonstrating that upon induction of TAD boundary pairing by light, paired alleles have a higher probability of being transcriptionally active We conclude that, as TADs are naturally dynamic, a snapshot of fixed cells at a given time may thus not reflect the actual relationships between chromatin structure and transcription activation. Our study highlights how this type of cause-effect relationship may be better uncovered by synthetic manipulative tools.

Our study has some limitations. First, our conclusions are based on observation of only two model TADs and genes, precluding us from drawing universally applicable conclusions regarding structure-function relationships of TADs. Second, while exploring the inherently dynamic process of TAD boundary pairing, we rely on data from fixed cells. The OptoTet system is fully suitable for live-cell imaging and future studies combining OptoTet with visualization of gene bursting in living cells should provide the necessary information to further dissect the dynamics and mechanisms underlying our observations.

Taken together, OptoTet is a synthetic biology approach to probe structure-function relationship of chromatin loops and domains. Its ability to manipulate chromatin structure in a controlled fashion on short time-scales and without the need for mutation of target sequences or depletion of proteins complements existing methods and should lead to novel insights into how chromatin structure relates to gene activity.

## ACKNOWLEDGMENTS

This research was funded by the Intramural Research Program of the NIH, NCI, Center for Cancer Research through grant 1-ZIA-BC010309 to TM and grant 1-ZIA-BC-011383 to DL and in part with Federal funds from the National Cancer Institute, National Institute of Health under Contract No. HHSN26120150003I. High-throughput imaging was performed in the NCI High-throughput Imaging facility (HiTIF) and confocal imaging was performed in the CCR/LRBGE Optical Microscopy Core, funded by the Intramural Research Program of the National Cancer Institute (NCI), Center for Cancer Research (CCR). This work utilized the computational resources of the NIH HPC Biowulf cluster (https://hpc.nih.gov). The contributions of the NIH author(s) were made as part of their official duties as NIH federal employees, are in compliance with agency policy requirements, and are considered Works of the United States Government. However, the findings and conclusions presented in this paper are those of the author(s) and do not necessarily reflect the views of the NIH or the U.S. Department of Health and Human Services.

## AUTHOR CONTRIBUTIONS

MS, TM conceptualized the project, designed the experiments, and interpreted the results. MS executed experiments. RC and VK, MP under supervision of DL, contributed to generation of cell lines. MS and TM drafted the manuscript. All authors participated in reviewing and editing the manuscript. TM, DL obtained funding.

## DATA AVAILABILITY

Output segmentation data is publicly available on Figshare (https://figshare.com/projects/mistelilab_optotet/283089). Scripts for analysis of segmentation data are publicly available on Zenodo (https://doi.org/10.5281/zenodo.23508428). Plasmids generated here are available in Addgene (Plasmid IDs #261184-261191).

