## Supplementary figures and tables for "Optogenetic induction of TAD boundary pairing promotes gene activity"

[2]Present Address: Instituto de Química Biológica de la Facultad de Ciencias Exactas y Naturales (IQUIBICEN), CONICET - Universidad de Buenos Aires, Facultad de Ciencias Exactas y Naturales, Buenos Aires C1428EGA, Argentina

### SUPPLEMENTARY FIGURES

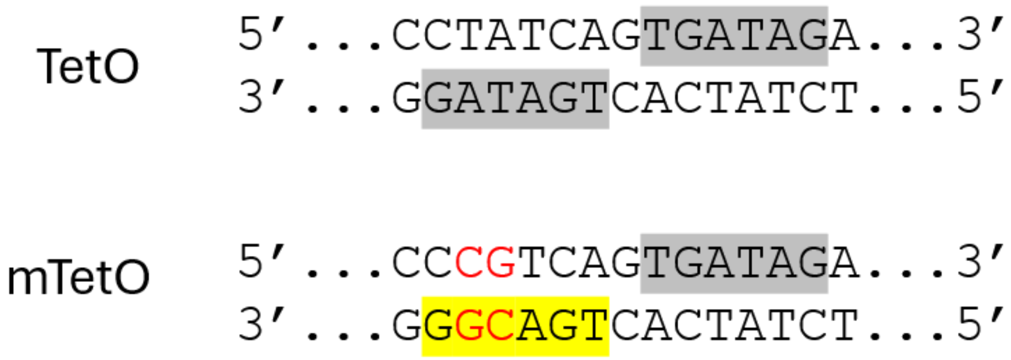


**Supplementary Fig. S1.** Sequence of TetO and mTetO repeats present in 96XTetO and 130XmTetO plasmids, respectively. Wild-type TetO is composed of two palindromic half-sites (highlighted in gray) each binding a wild-type TetR monomer. Mutated TetO (mTetO) is composed of a wild-type half-site and a mutated half-site (highlighted in yellow). Mutated base pairs are marked in red.

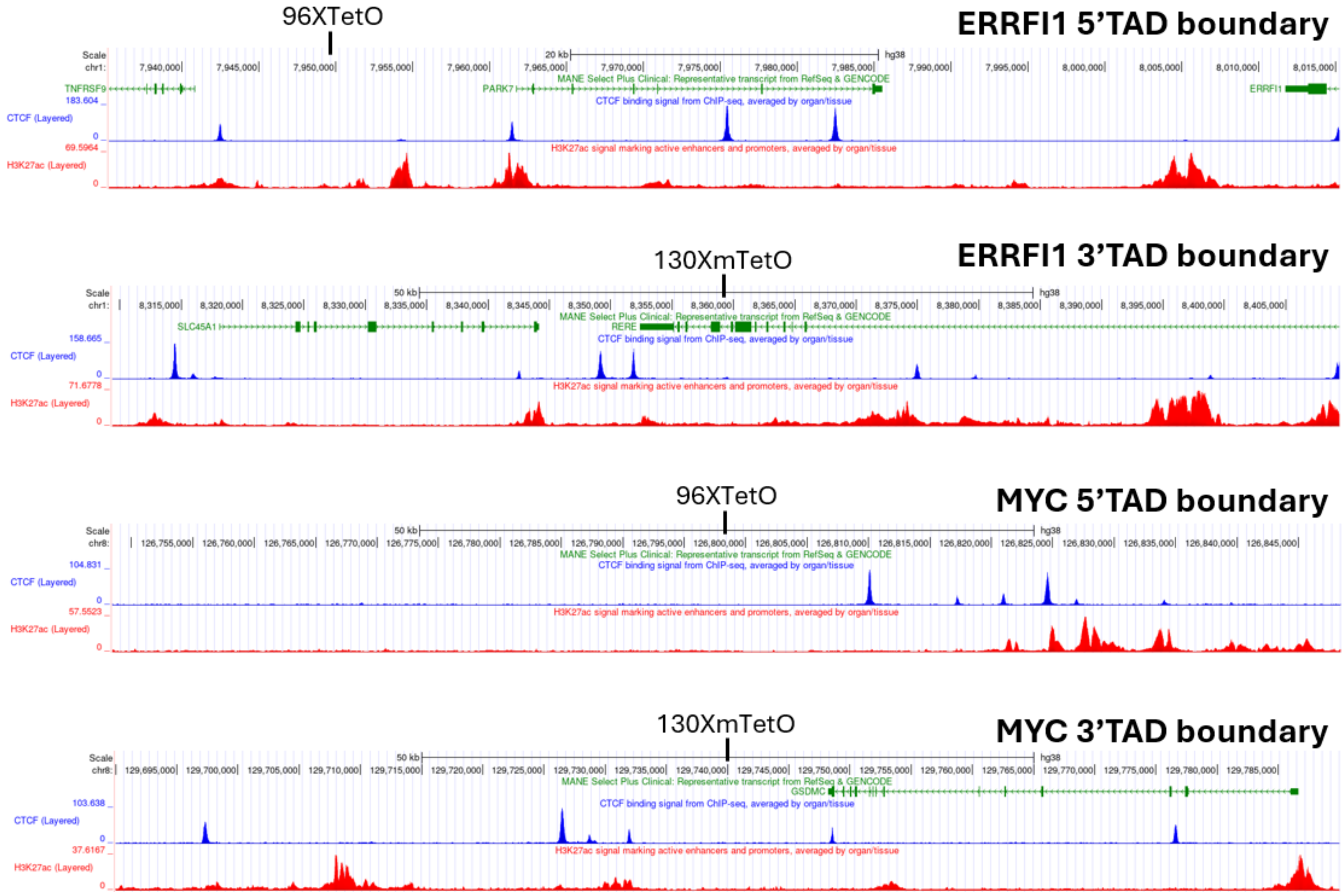


**Supplementary Fig. S2.** Genome Browser (https://genome.ucsc.edu) showing *ERRFI1* and *MYC* TAD boundaries with their surrounding regions, with tracks representing MANE representative transcripts from RefSeq and GENCODE (green), CTCF ChIP-seq data (blue, averaged from 51 tissues/organs from ENCODE 4), and H3K27ac ChIP-seq data (red, averaged from 55 tissues/organs from ENCODE 4). Locations of integrations of TetO arrays are indicated.

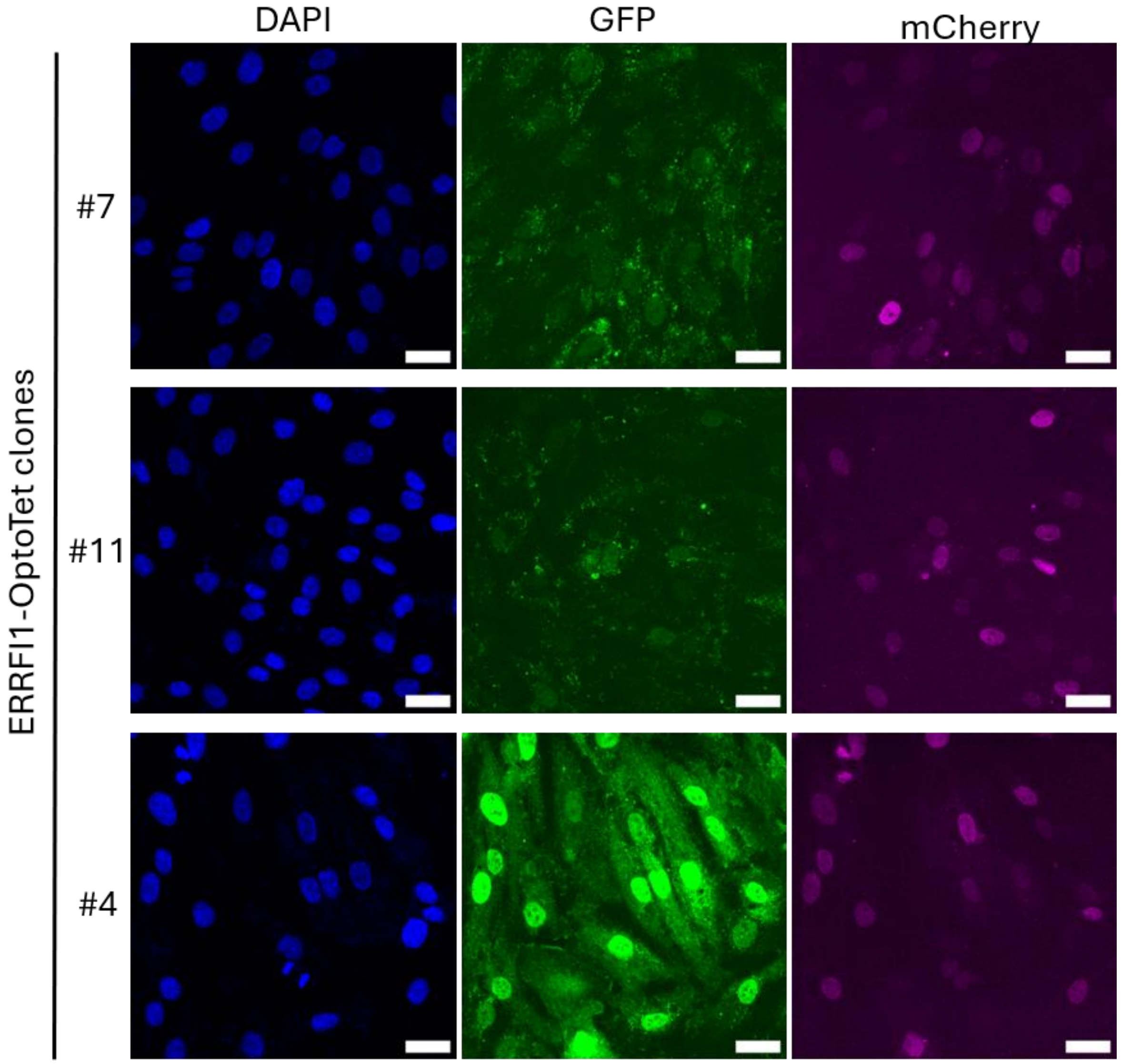


**Supplementary Fig. S3.** Representative images of ERRFI1-OptoTet clones with stable expression of TetR-GFP-CRY2 and mTetR-mCherry-CRY2. The same intensity scales were set for all images to compare expression levels between clones. Scale bars: 30 μm.

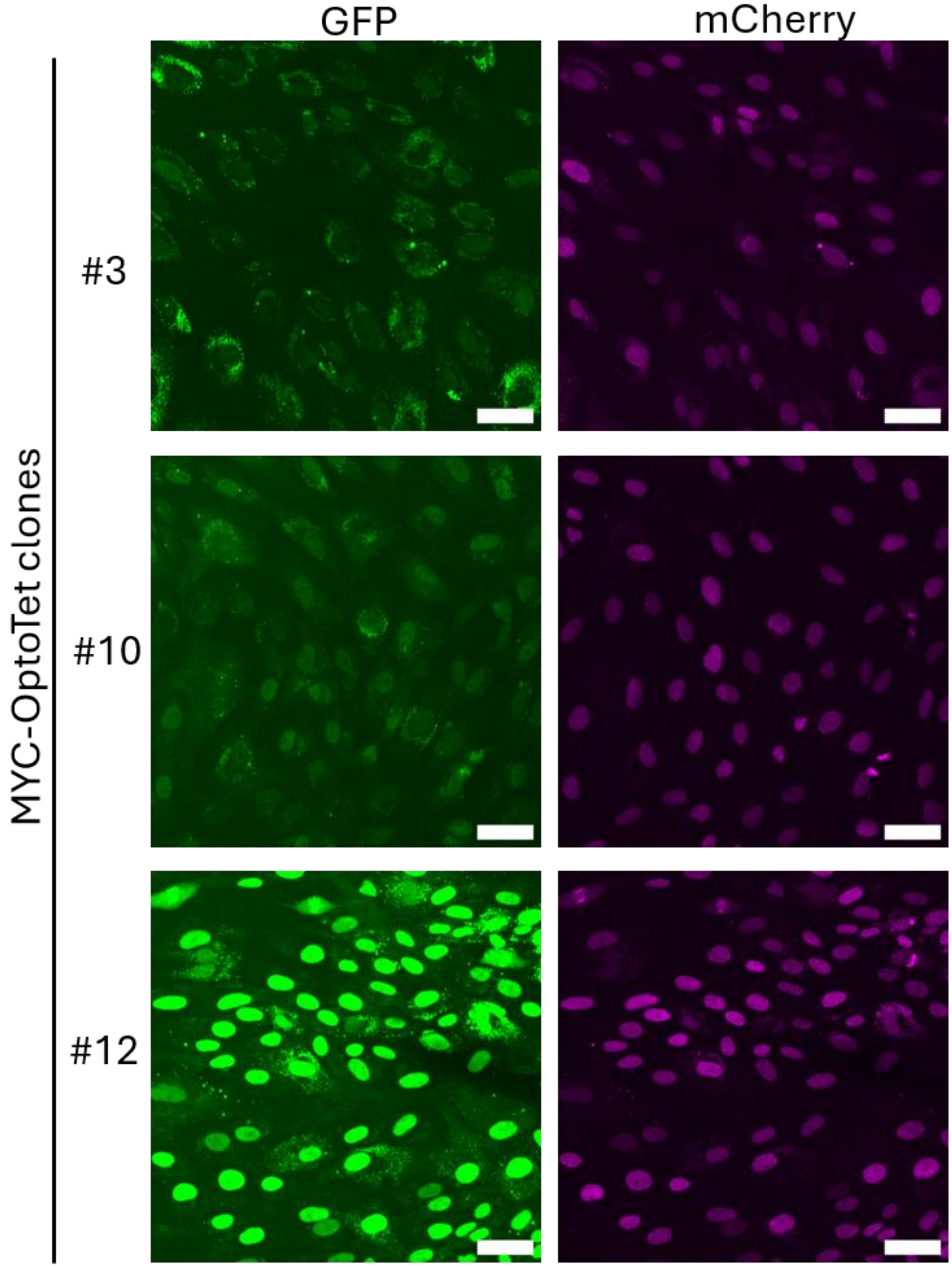


**Supplementary Fig. S4.** Representative images of MYC-OptoTet clones with stable expression of TetR-GFP-CRY2 and mTetR-mCherry-CRY2. The same intensity scales were set for all images to compare expression levels between clones. Scale bars: 50 μm.

**SUPPLEMENTARY TABLES**

**Supplementary Table S1. List of plasmids**

| Plasmid | Source | Reference |
| --- | --- | --- |
| ERRFI1-5pTAD-96XTetO | Generated in this study | Deposited in Addgene (#261184) |
| MYC-5pTAD-96XTetO | Generated in this study | Deposited in Addgene (#261185) |
| ERRFI1-3pTAD-130XmTetO | Generated in this study | Deposited in Addgene (#261186) |
| MYC-3pTAD-130XmTetO | Generated in this study | Deposited in Addgene (#261187) |
| pHAGE_scTetR-3XGFP | Generated in this study | Deposited in Addgene (#261188) |
| pHAGE_TetR-mTetR-3XmCh | Generated in this study | Deposited in Addgene (#261189) |
| pHAGE_scTetR-3XGFP-CRY2 | Generated in this study | Deposited in Addgene (#261190) |
| pHAGE_TetR-mTetR-3XmCh-CRY2 | Generated in this study | Deposited in Addgene (#261191) |
| psPAX2 | Addgene #12260 | Generated by Didier Trono |
| pMD2.G | Addgene #12259 | Generated by Didier Trono |

**Supplementary Table S2. List of single-guide RNAs.**

| Name | Spacer sequence | Source | Use |
|---|---|---|---|
| 5P-ERFFI1-IVT-4698 | GUUAAGGUGAACGCUAGGGA | Synthego | 96XTetO knock-in at ERRFI1 5'TAD boundary |
| 3P-ERRFI1-IVT-4727 | CAGGCGAUAAGCAGGCUGCG | Synthego | 130XmTetO knock-in at ERRFI1 3'TAD boundary |
| 5P-MYC-IVT-4732 | CGUUGUGCCUAGAACCACUA | Synthego | 96XTetO knock-in at MYC 5'TAD boundary |
| 3P-MYC-IVT-4760 | CAUUAGCCCAGUCCUACGUA | Synthego | 130XmTetO knock-in at MYC 3'TAD boundary |
| sgRAD21_1 | CAUUUUGUUCUCAGUAAAAG | EditCo | RAD21 knockout |
| sgRAD21_2 | GUCAACAUCAAACAUACCUU | EditCo | RAD21 knockout |
| sgRAD21_3 | ACACUCGAACACAUGGGCUU | EditCo | RAD21 knockout |
| sgNIPBL_1 | UAUAAUUAGUUCUCCAUCCA | EditCo | NIPBL knockout |
| sgNIPBL_2 | UCGUGUAAGGUCUUCAGACA | EditCo | NIPBL knockout |
| sgNIPBL_3 | CCUUUUAAUGUGCAGUACCC | EditCo | NIPBL knockout |

**Supplementary Table S3. List of oligonucleotide primers for PCR**

| Name | Sequence | Use |
|---|---|---|
| Zeo_Fw | CAAGTTGACCAGTGCCGTTCCG | Screen 5'TAD insertions |
| ERRFI1_5p_Rv | GACACTGGTCAAAGAATGCTG | Screen ERRFI1 5'TAD insertion |
| MYC_5p_Rv | AACACTGAGTGCCTCTTGACTG | Screen MYC 5'TAD insertion |
| Neo_Fw1 | CTATCGCCTTCTTGACGAGTTCTTCTGAG | Screen ERRFI1 3'TAD insertion |
| ERRFI1_3p_Rv | CCGTTTGCTCAGCACCCCTTTGTC | Screen ERRFI1 3'TAD insertion |
| Neo_Fw2 | GTTATCTCGAGAGCGCTGCTAGCTTAAG | Screen MYC 3'TAD insertion |
| MYC_3p_Rv | CTCCATGACACTGGTCTGGGCAATG | Screen MYC 3'TAD insertion |

**Supplementary Table S4. List of reagents for cell line knock-in**

| Location (hg38) | Target cell line | Donor DNA | sgRNA | Primers for screen | Final cell line product |
|---|---|---|---|---|---|
| chr1: 7949608 | hTERT-RPE1 | ERRFI1-5pTAD-96XTetO | 5P-ERFFI1-IVT-4698 | Zeo_Fw + ERRFI1_5p_Rv | ERRFI1-5'TAD |
| chr8: 126798442 | hTERT-RPE1 | MYC-5pTAD-96XTetO | 5P-MYC-IVT-4732 | Zeo_Fw + MYC_5p_Rv | MYC-5'TAD |
| chr1: 8359383 | ERRFI1-5'TAD | ERRFI1-3pTAD-130XmTetO | 3P-ERRFI1-IVT-4727 | Neo_Fw1 + ERRFI1_3p_Rv | ERRFI1-5'/3'TAD |
| chr8: 129739993 | MYC-5'TAD | MYC-3pTAD-130XmTetO | 3P-MYC-IVT-4760 | Neo_Fw2 + MYC_3p_Rv | MYC-5'/3'TAD |

**Supplementary Table S5. List of stable cell lines with FP constructs**

| Name | Stable expression | Lentiviral constructs |
|---|---|---|
| TetR alone | TetR-GFP<br>mTetR-mCh | pHAGE_scTetR-3XGFP<br>pHAGE_TetR-mTetR-3XmCh |
| OptoTet | TetR-GFP-CRY2<br>mTetR-mCh-CRY2 | pHAGE_scTetR-3XGFP-CRY2<br>pHAGE_TetR-mTetR-3XmCh-CRY2 |

**Supplementary Table S6. List of BAC probes for DNA-FISH (Empire Genomics).**

| BAC | Target | Dye | Location (hg38) |
|---|---|---|---|
| RP11-767K22 | ERRFI1 5' TAD boundary | 'Orange' | chr1:7880719-8065702 |
| RP11-366K21 | ERRFI1 3' TAD boundary | 'Green' | chr1:8241991-8425192 |
| RP11-765K23 | MYC 5' TAD boundary | 'Red' | chr8:126761149-126937724 |
| RP11-717D13 | MYC 3' TAD boundary | 'Green' | chr8:129618087-129782546 |

**Supplementary Table S7. List of oligonucleotide probes for *ERRFI1* RNA-FISH.**

Designed as a pool and labeled with Atto647 ( LGC Biosearch Technologies).

| **Sequence** |
|---|
| ATGTGACACATTTGCTGTGG |
| AGTAAGGGATCAGTGCAAAT |
| CTATTGTGTAACACTGACCC |
| GCACCAGGATATTATTTGTC |
| AAGTTTAGGTTTTTTTCCTT |
| TGTCTTCAGATAGCCAAGTT |
| GTTAGTCAGCATCAATCTCT |
| TAGTTCCTCAACTTCAGGAC |
| AGCTGCAAGGCATGGATAAC |
| CACCCAGTAACTCATATCTT |
| GGTACATTCTTTACATCCTC |
| TCAGTAAAAGCCTGGGACAG |
| AAGTTCTGCAAGATTACTGG |
| ACTAATTTCCATTATCCTGT |
| TTCTCCAGTTACTACTTGTT |
| GAGTTTCTACAGTATTGGCA |
| AGCCACTGTATTTTATGCTG |
| CCTGACTTATTAAACCAGCC |
| CAAAAGCAGACCTCCTACTG |
| CGTGAGGGTACTTATATGTA |
| CTAACAGAGGATTTGGGGGA |
| CCTCCATTAGGACAGTCTAA |
